# High-throughput assessment of exercise-induced adaptations and muscle function in health and ageing

**DOI:** 10.1101/2025.01.16.633393

**Authors:** Alina Malita, Michael J. Texada, Olga Kubrak, Mette Lassen, Anne H. Skakkebaek, Ivan Bradic, Martin R. Larsen, Nadja Ahrentløv, Kim Rewitz

## Abstract

Exercise improves metabolic health, enhances insulin sensitivity, and preserves muscle function, making it a core intervention to combat age-related decline and metabolic disorders. However, large-scale genetic and pharmacological screens to uncover exercise-induced adaptations remain challenging in mammalian models due to their complexity and cost. Here, we present the ClimbMaster, a fully automated, computer-controlled platform for assessing exercise-induced adaptations and physical performance in the fruit fly *Drosophila*. The system uses repeated climbing exercises to simulate endurance training, enabling precise measurements of climbing speed and endurance across different conditions and life stages. Using the ClimbMaster, we demonstrate that exercise improves endurance, promotes fat loss, and enhances insulin-mediated glucose uptake and modulates insulin sensitivity in muscle, highlighting key conserved features of exercise physiology between flies and mammals. We also show that rapamycin, a TOR inhibitor, mitigates age-related performance decline. This platform enables high-throughput screens to investigate genetic and environmental factors influencing muscle health, aging, and insulin resistance, providing a scalable and versatile tool for the identification of novel therapeutic targets to improve healthspan and physical performance.

## Introduction

Physical activity has long been recognized as a powerful tool for improving health and preventing chronic diseases^1^. Regular exercise leads to a range of health benefits, including enhanced cardiovascular fitness, improved glucose metabolism, and reduced risk of metabolic disorders^2^. However, muscle mass and strength naturally decline with age, contributing to frailty, reduced mobility, and an increased risk of metabolic dysregulation^3^. Maintaining muscle mass is therefore critical for healthy aging and for mitigating the effects of age-related decline^4^. Furthermore, it has become clear that recent breakthroughs in obesity treatments, such as GLP-1 receptor agonists^5^, while highly effective, can lead to significant unintended muscle loss^6^. This loss of muscle mass poses a serious risk, as reduced muscle mass is associated with impaired physical function, slower metabolism, and an increased risk of falls, fractures, and metabolic diseases^7^.

Skeletal muscle plays a central role in maintaining whole-body metabolism^8^. Acting as a major sink for glucose, muscle tissue helps regulate blood glucose levels and burns a substantial portion of the body’s energy stores^9,10^. Therefore, increasing muscle mass can improve metabolic health, enhance insulin sensitivity, and reduce the risk of type-2 diabetes (T2D). Anabolic steroids can promote muscle growth, but their use is limited due to severe side effects, including cardiovascular risks and hormonal imbalances^11^. This underscores the urgent need to identify non-androgenic strategies that preserve or increase muscle mass, both to counteract age-related decline and to mitigate muscle loss during weight-management therapies. Across a wide spectrum of human diseases, strategies that prevent muscle loss may have the greatest impact on improving diseases, healthspan and quality of life; such interventions have therefore become a major focus of current research.

As a means to identify new ways of preserving muscle, there is an increasing demand for *in-vivo* models that provide mechanistic insights into the regulation of muscle mass and strength, as well as platforms for large-scale drug screening to discover novel non-androgenic anabolic factors. While *in-vitro* systems are useful for studying individual pathways, they fail to capture the complexity of whole-organism responses, including inter-tissue communication and the impact of metabolism, hormonal regulation, and neural inputs on muscle function. *In-vivo* models are essential for identifying systemic factors that influence muscle health and for evaluating potential interventions in a physiologically relevant context^12^. Mammalian models, such as mice, are widely used for studying muscle biology. However, large-scale genetic screens and drug-discovery efforts are impractical in mammalian models due to high costs, long lifespans, and ethical considerations. The complexity and expense of maintaining mammalian colonies, combined with the time required to assess muscle phenotypes over the lifespan, make performing high-throughput studies in these systems a challenging endeavor. Furthermore, exercise protocols in mammals are labor-intensive and often require specialized equipment, which limits their scalability for large studies.

The fruit fly *Drosophila* offers a unique and powerful *in-vivo* system for studying muscle function and metabolism^13^. The fruit fly shares conserved insulin signaling pathways, which are central to metabolic regulation, and has a well-characterized genetic toolkit that allows for precise manipulation of gene function^14,15^. Remarkably, the fly insulin pathway is so highly conserved that it can be activated by human insulin^16^. Human GLUT4 – the transporter that carries out insulin-stimulated glucose uptake in the fat and musculature – expressed in the fly recapitulates mammalian glucose-uptake regulatory mechanisms, exhibiting insulin-induced translocation to the plasma membrane^17^. This high degree of conservation makes the fly a particularly relevant model for studying insulin-regulated muscle metabolism. *Drosophila* thus offers a unique, powerful system for studying muscle function and metabolism. Systems that exploit flies’ innate negative geotaxis behavior (their instinct to climb upward after being displaced downward) by repeatedly tapping the animals down have been shown to induce exercise effectively^18^. This approach can also be used to assess the animals’ physical performance, specifically their climbing ability, which is a well-established proxy for muscle strength and endurance^18,19^. However, despite these advantages, a high-throughput platform for implementing robust and precise exercise regimes paired with automated assays to monitor physical performance has been lacking in flies.

To address this need, we developed a fully automated computer-controlled system that can precisely administer exercise protocols and automatically track climbing performance in *Drosophila*, which we term the ClimbMaster. This apparatus provides a reliable, high-throughput, and precise method to assess physical performance in the fly by measuring both climbing speed and endurance during repeated exercise sessions, as well as over time during aging. Here, we present a comprehensive framework for implementing robust exercise regimes in flies and tracking their performance over time. We use this system to demonstrate sexual dimorphism in physical endurance, and we show that exercise regimes improve physical performance, promote healthy sleep and activity patterns, make flies leaner, and reveal performance declines during aging. Furthermore, we demonstrate that exercise affects the molecular physiology of muscle tissue, enhancing glucose uptake and modulating insulin responsiveness, indicating that the ClimbMaster can be used to study the systemic effects of physical activity. We also show that treatment with the TOR inhibitor rapamycin increased performance in aging animals, highlighting the system’s potential for identifying drugs that prevent age-related performance decline. These findings suggest that the ClimbMaster provides a versatile platform to investigate the benefits of exercise across multiple biological scales, from behavioral outputs to molecular pathways. The broad range of effects observed highlights the wide applicability and translational potential of this model for studying exercise physiology and muscle health, with implications for both basic research and therapeutic development. Ultimately, the system has the potential to drive new discoveries that contribute to better strategies for preserving muscle health and improving quality of life across the lifespan.

## Results

### Development of the ClimbMaster

To study the physiological and molecular adaptations to endurance exercise in *Drosophila melanogaster*, we developed the ClimbMaster, a custom-built, automated apparatus (Fig. 1a). The system leverages flies’ innate negative geotaxis behavior, in which they naturally climb upward after being displaced downward. The ClimbMaster consists of a modular design with an automated displacement mechanism powered by digital servos, which tap flies downward at precisely timed intervals. Each vial, containing ∼20 flies, is securely positioned in an 8 x 6 grid, allowing simultaneous exercise of up to 48 experimental groups. Of these, 16 vials (2 x 8 on each side) can be monitored in real time through video recording, enabling precise tracking of climbing performance. The system is operated via an advanced controller box connected to a computer running custom-designed software, enabling standardized, high-throughput exercise protocols. This setup enables precise programming of displacement intervals, exercise duration, and experimental conditions, ensuring reproducibility and flexibility for a wide range of study designs. The integration of automated hardware with custom code provides researchers with a powerful tool for standardized, high-throughput exercise experiments. The typical exercise protocol involves one hour of repeated downward displacements at 8-second intervals. This system, much like treadmills or wheel-running setups in mammalian exercise studies, provides a controlled and reproducible framework for inducing repetitive physical activity. By enabling the study of endurance and its physiological consequences, the ClimbMaster serves as an analogous platform to investigate conserved molecular and physiological responses to exercise across species.

**Figure 1.**
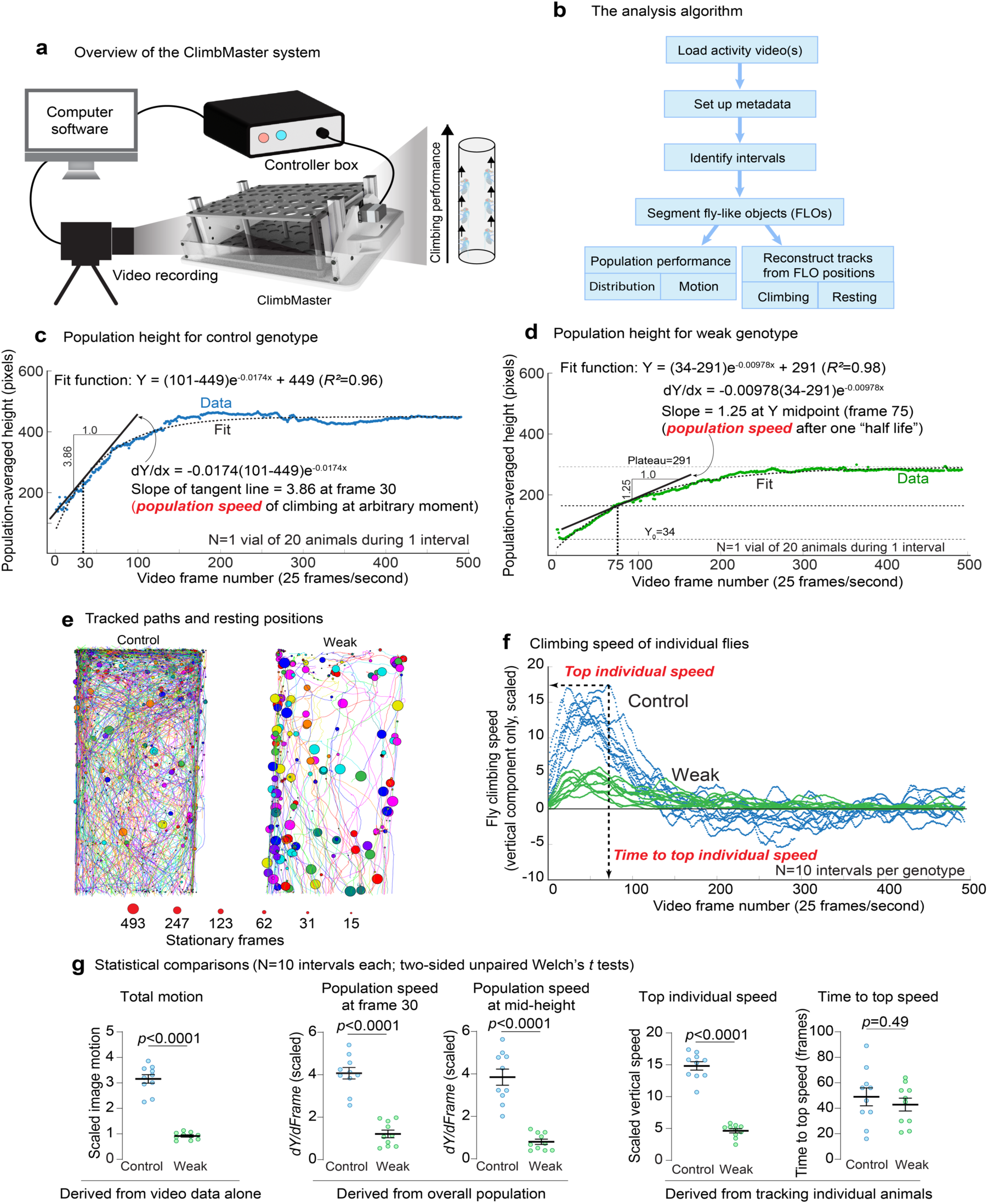
Design and performance output of the ClimbMaster exercise apparatus. (a) The ClimbMaster is a custom-built, automated device for studying endurance exercise in fruit flies. It features a modular vial grid for housing multiple groups of flies, with each vial holding 20 flies. Digital servos displace vials downward every 8 seconds, triggering negative geotaxis. The system integrates automated hardware with custom software for standardized, high-throughput exercise experiments. (b) A flowchart of the processing algorithm. (c) The “center of mass” of the population of detected flies is shown over time for one vial of ∼20 control flies during one interval of recording (magenta points). A one-phase exponential-decay curve is fit to this performance data (dotted black line). The first derivative of this function gives the speed of movement *of the population midpoint* at the arbitrarily chosen time point frame 30; this is referred to elsewhere as the population speed. (d) Similar to (c), for flies of a weak genotype with a muscle-specific mitochondrial defect (green) tested simultaneously. The slope of the fit function – the speed of the population as a whole – is also taken at time point when the population reaches the midpoint between the minimum and plateau heights. This time point reflects one “half life” of the decay function and thus is not fixed at an arbitrary video frame but instead varies from assay to assay. (e) Reconstructed tracks of flies in the ten performance intervals for the two genotypes in (b) and (c). Each reconstructed track is indicated by a line, and the last position along the track is indicated by a circular marker. Larger dots indicate that the animal had been at that position for a longer time when the track ended. Note that the weak genotype gives a much sparser pattern of tracks, with the animals reaching a resting position sooner and remaining there for a longer time. (f) The climbing speed of individually tracked flies, averaged across all the flies in each vial, during each of ten intervals. The curves here are smoothed using a running average. The average speed increases rapidly after each “banging” event, perhaps as the animals spread out and can move freely, after which their speed begins to decrease (perhaps as they begin to crowd one another at the top of the vial) and can even turn negative (as the animals move away from the top of the vial to spread out from one another). For each test interval, the peak value of the running-averaged speed is recorded, along with the video-frame number at which that peak was reached. (g) Statistical comparisons of the derived performance values illustrated in the panels above. Panel #1: the number of pixels that differ from one frame to the next are summed for each interval and each vial, as a simple measure of motion that does not rely on tracking. Panel #2 and #3: the slopes of the fit curves for population height over time – that is, the population speed – during each interval at the arbitrarily chosen fixed time Frame 30 (#2) and at the variable time that the Y midpoint is reached (#3). Panel #4 and #5: the peaks of the speed curves shown in (f) and the time at which these peaks occur. All comparisons are two-sided unpaired Welch’s *t* tests.

### Development of an automated system for quantifying climbing performance in *Drosophila*

To measure climbing performance, flies were video recorded during 10 consecutive rounds of downward displacement, with 20-second intervals between rounds to allow time for a full behavioral response. Videos were analyzed using custom-written software to quantify climbing speed, endurance, and consistency. Existing analysis packages^20^ were not suited to the repeated-test nature of these assays, required video preprocessing, or were not designed to produce the desired analytical outputs. To meet our needs, we therefore developed an analysis system in the MATLAB environment (MathWorks, Natick, MA), specifically designed to analyze fly climbing performance in video recordings of repeated behavioral intervals. The script interactively processes single videos or, in batch mode, multiple videos without user interaction after the initial setup. It detects the displacement events that demarcate the repeated analysis intervals, and for each of these intervals, the code identifies fly-like objects within each vial, reconstructs their paths of locomotion, and generates numerical outputs reflecting speed and position data (Fig. 1b). These analyses generate comprehensive readouts of climbing performance for each vial during each test interval. The script requires the MATLAB Image Processing, Computer Vision, and Curve Fitting toolboxes and will be made freely available upon request.

The script analyzes climbing performance in two complementary ways: (1) by observing the “center of mass” of all identified flies as a group over time and (2) by tracking individual flies and reconstructing their movements over time. To ground-test the system, we video-recorded control animals and animals in which we induced defects in muscle mitochondrial function, to be described separately, over ten bouts of exercise. As an indicator of total activity during each interval, the differences between consecutive frames (pixels that changed) were summed into a readout of “total motion” (Fig. 1g, left). Control animals exhibited greater activity in this readout compared to animals with muscle mitochondrial defects, referred to as weak genotypes hereafter (*p*<0.0001).

The “center of mass” of the animals – the average height of the flies within each vial, during each bout (“interval”) – rose over time after the tap-down stimulus and reached a steady state as the animals attained a distribution that satisfies their innate rules for spacing between animals and other objects (Fig. 1c for one interval of control animals and 1d for the weak genotype). Note that the weak genotype settles into a lower steady state than the controls, and these animals take a longer time to reach this state. The curve of the position of the population center over time can be fit by a single-phase exponential-decay function (*R^2^* in Fig. 1c, 0.96; in Fig. 1d, 0.98), and the slope of this line at any point reflects the instantaneous speed at which the population is moving. We compared these slopes at the fixed arbitrary position of frame 30 in each interval, which corresponds to 1.2 seconds after the downward displacement (Fig. 1g, second panel), and at the frame where each curve reaches the midway point between the minimum and maximum heights (Fig. 1g, third panel), a time point that is mathematically characteristic of each curve (one “half life”). In both cases, the weak genotype displayed less robust performance than the control (*p*<0.0001).

The script also identifies and tracks single flies over time as a second route of analysis. Individual reconstructed tracks for the control and weak genotypes (Fig. 1e) are shown with a marker at each animal’s last position, the size of which indicates the length of time the animal had been in that position when the interval ended – animals that stopped moving earlier in the interval, such as those of the weak genotype here, are indicated by larger markers. The vertical component of all of the individual flies’ locomotion is averaged at each frame, and these are smoothed using a running average to give a curve of the average individual speed for each vial during each interval (Fig. 1f). The animals’ speed (on average) increased after the downward displacement, perhaps reflecting first disorganized motion as the animals interfere with one another, followed by faster movement as the animals spread out. The average speed reaches a peak before falling again, possibly as the animals begin crowding into the top of the vial; negative speeds indicate that the animals begin moving downwards again, potentially to attain more “comfortable” spacing. The peak value in each assay, and the time point at which this peak is reached, are useful data for comparison (Fig. 1g, panels 4 and 5). The animals of the control genotype reached a higher average speed than those of the weak strain (*p*<0.0001), but they reach this speed with the same latency. These readouts provide a comprehensive description of the animals’ climbing performance in the ClimbMaster, and thus they permit the investigation of a number of different aspects of the animals’ behavior.

### Sex-specific differences in climbing endurance

We explored potential sex differences in physical climbing performance by subjecting male and female animals to a prolonged exercise regimen in the ClimbMaster platform. Animals with no prior exercise training were exercised in a 4.5-hour climbing routine with 8-second tap-down intervals, and their performance was measured at 30-minute intervals during the routine. Initial measurements prior to exercise revealed few differences in climbing performance between sexes, as both males and females displayed similar metrics for all the parameters evaluated (0-h timepoints in Fig. 2a–d). The top individual speed (*i.e.*, the speed of individually tracked animals averaged at each frame in the video), a measure of peak performance, initially increased from the initial assessment to a maximum at the observation after 30 minutes of exercise. This early effect of stimulation, perhaps due to stimulus-driven excitation of neural or muscular tone, suggests that both initial performance and performance after warming up might be usefully explored.

**Figure 2.**
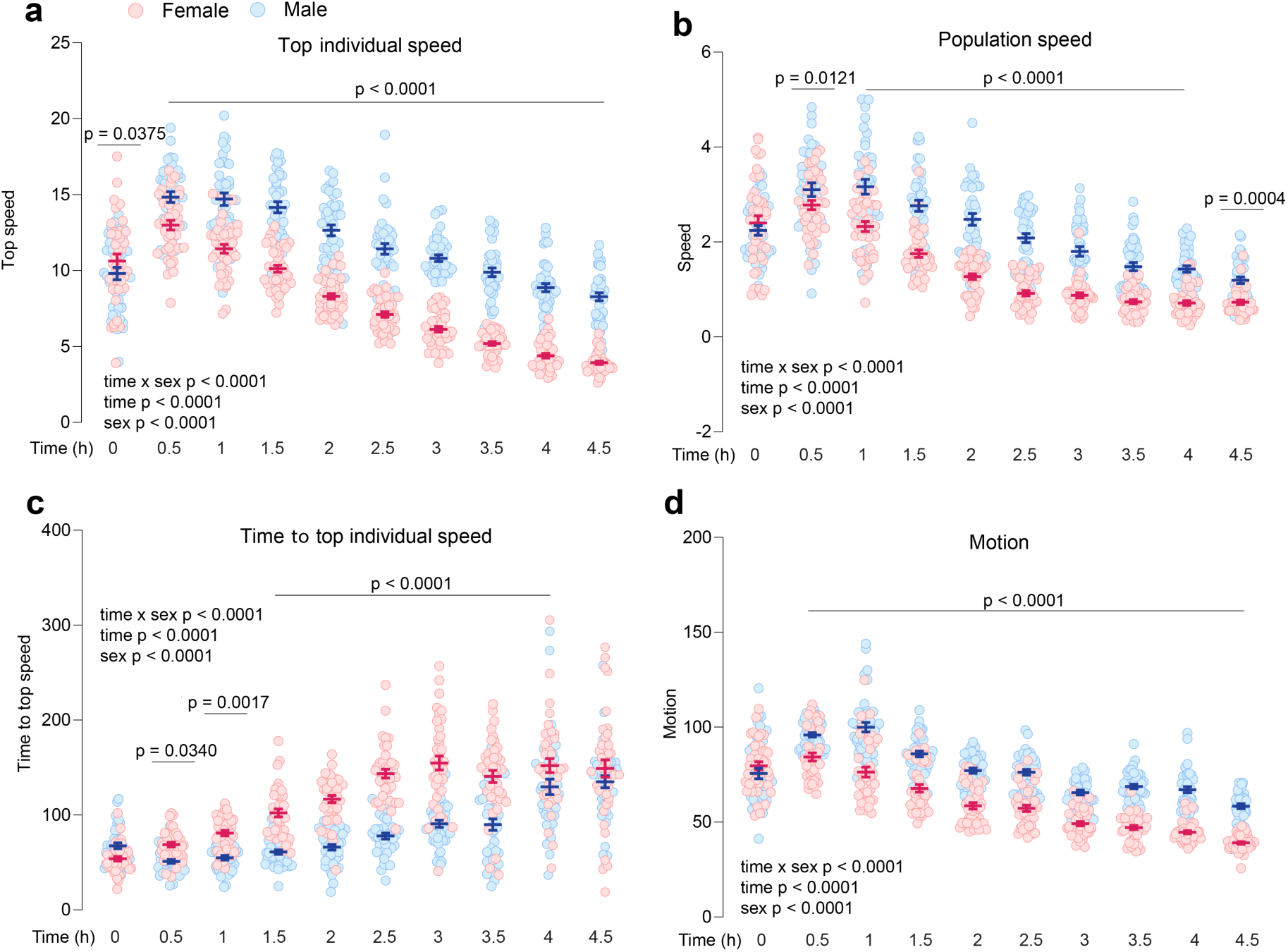
Sexual dimorphism in climbing performance. Climbing performance was measured in male and female flies subjected to a 4.5-hour exercise regimen with 8-second tap-down intervals using the ClimbMaster platform. Performance was recorded at 30-minute intervals, and multiple climbing metrics were analyzed. (a) “Top individual speed” represents the highest average climbing speed of individually tracked animals at each time point, and (b) “time to top individual speed” reflects the time required for flies to reach their maximum speed during each interval, as in Fig. 1f. (c) “Population speed” represents the speed at the Y-achieved by the group of animals at each time point, as in Fig. 1d, while (d) “motion” captures overall movement during the exercise period. Data are presented as individual data points with mean ± s.e.m. Statistical significance was determined using two-way ANOVA, with significant effects of sex, time, and sex × time interactions for all performance metrics (*p* < 0.0001). Statistical significance of the male-vs.-female difference at each time point was determined by Tukey’s multiple-comparisons test.

Sexual dimorphism in performance emerged over the length of the exercise routine. Males consistently exhibited higher top individual speeds than females, with the difference becoming more pronounced over time as female speeds dropped more quickly, suggestive of faster tiring and exhaustion in these animals (Fig. 2a). The time required to reach the peak in averaged individually tracked speed increased progressively in both sexes as the exercise regimen advanced, suggestive of fatigue. However, females displayed a more substantial increase in this parameter earlier in the exercise regimen compared to males (Fig. 2c). These results suggest that males experienced a slower decline in physical ability over time compared to females and were less affected by the cumulative effects of fatigue. In the independently derived population measurement, which does not rely on tracking individual animals and thus reflects the climbing performance of the animals as a group, males consistently outperformed females across the entire exercise period after the initial assessment. This second method supports the individual-animal tracking and the existence of sexual dimorphism in performance under sustained exertion (Fig. 2b). Finally, motion, a parameter capturing the overall activity level of the animals (that is, reflecting any movement, not just vertical locomotion), showed trends mirroring top speed, with males maintaining higher levels of activity compared to females during prolonged periods of intense physical activity (Fig. 2d). Performance differences over time and between sexes were statistically strong (two-way ANOVA factor effects *p*<0.0001 for both), and significant interaction between sex and time was detected for all performance parameters (two-way ANOVA sex × time interaction *p*<0.0001), indicating that the trajectory of performance changes differed between males and females.

Together, these results reveal significant sex differences in climbing performance in *Drosophila*, with males exhibiting prolonged endurance and peak performance during sustained exercise. These findings align with observations in mammals, in which males often display greater endurance and physical performance. The interaction between sex and time underscores the importance of considering sex as a biological variable in studies of physical performance and endurance. The biological drivers of these differences in the fly will be fruitful topics of future exploration.

### Climbing performance declines with age and is impaired by a high-sugar diet whereas rapamycin treatment has beneficial effects in older flies

To investigate the effects of aging and diet on climbing performance, we tested male and female flies at multiple time points and under different dietary conditions using the ClimbMaster platform. Climbing performance was assessed at one week, two weeks, and four weeks of age to evaluate age-related changes. Aging significantly impaired physical performance in both males and females (Fig. 3a, b). Top individual speed, a measure of peak climbing performance, declined progressively with age in both sexes (Fig. 3a). Similarly, the time to reach top speed increased with age, indicating slower responses and reduced physical endurance in aged flies (Fig. 3b). These findings demonstrate that *Drosophila* exhibit age-related declines in physical performance similar to those observed in other animal models, highlighting the utility of this climbing assay for studying age-associated physiological changes.

**Figure 3.**
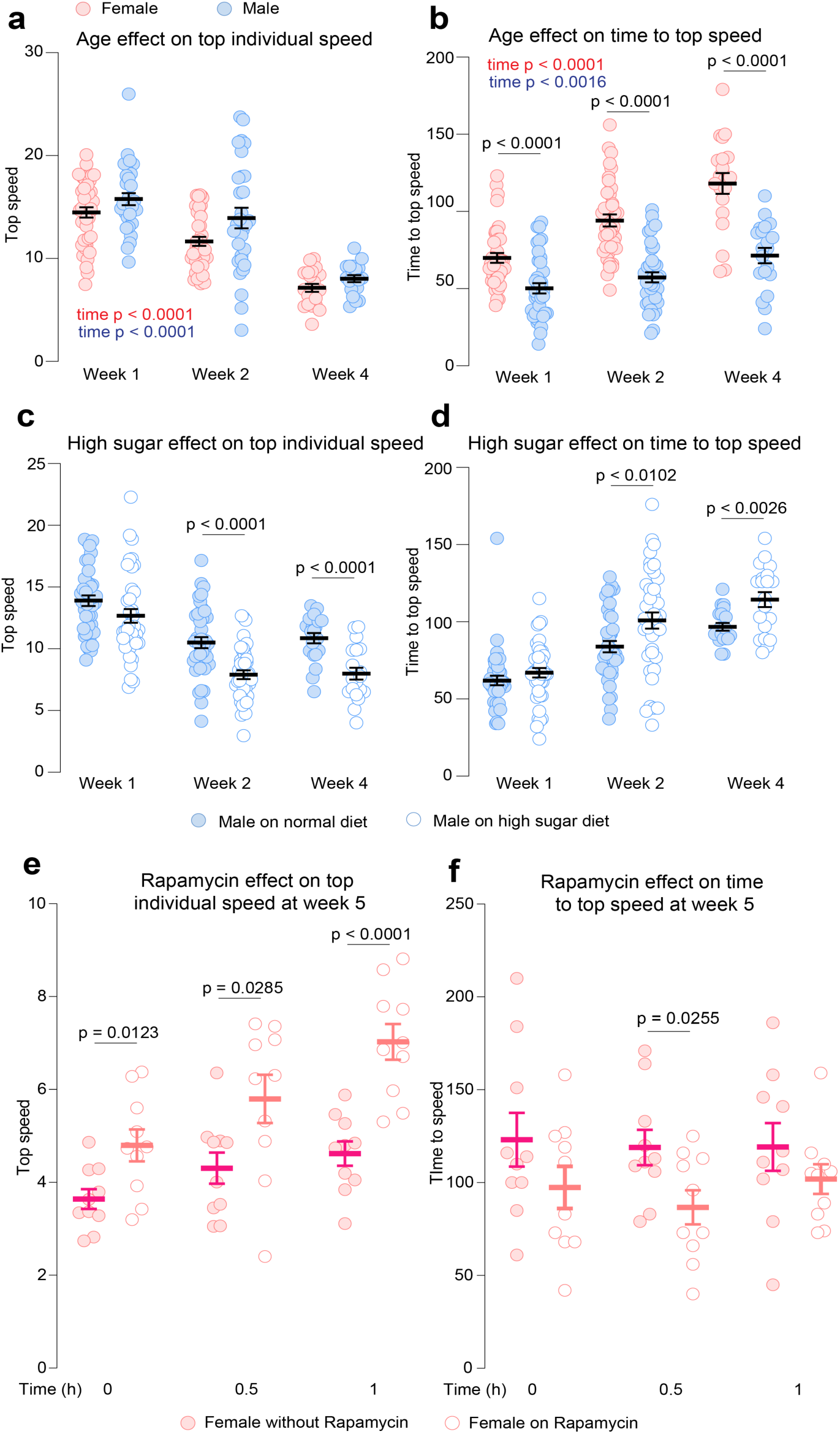
Aging and high-sugar diet impair climbing performance whereas inhibition of mTOR increases physical performance in older animals. Climbing performance was measured in male and female flies using the ClimbMaster platform at one week, two weeks, and four weeks of age. (a) Top individual speed in male and female flies across the three time points. (b) Time to reach top speed in male and female flies across the same time points. (c) Top individual speed in male flies fed a normal diet (8% sugar) or a high-sugar diet (20% sugar) across the three time points. (d) Time to reach top speed in male flies on normal and high-sugar diets across the same time points. (e) Top individual speed and (f) Time to reach top speed in 5-week-old females fed a normal diet or supplemented with rapamycin before, 30-minutes and 1-hour during the exercise. Data are presented as individual data points with mean ± s.e.m. Statistical significance was determined using one-way ANOVA for panels (a) and (b) and with two-tailed *t-*tests for panels (c), (d), (e) and (f).

We next tested the effect of a high-sugar diet, which mimics obesity and diabetic phenotypes in flies^16,21,22^, on age-related climbing performance. Male flies were maintained on either a normal diet containing 9% sugar or a high-sugar diet (HSD) containing 20% sugar, with yeast as the primary protein/lipid source kept constant. Flies fed HSD showed impaired climbing performance compared to those fed a normal diet (Fig. 3c, d). Males on a high-sugar diet exhibited significantly lower top individual speeds at the two-week and four-week time points (Fig. 3c) and took longer to reach their top speed at those same time points (Fig. 3d), suggesting that a high-sugar diet exacerbates the decline in physical performance with age. These results indicate that the high-sugar dietary model is a relevant tool for studying metabolic dysfunction and its impact on physical performance during ageing.

During aging, processes such as muscle protein synthesis and autophagy become dysregulated, leading to physical decline^23,24^. Since these processes are regulated by the TOR pathway, we aimed to determine whether inhibiting the TOR pathway through rapamycin treatment could enhance physical performance in older flies. We found that climbing performance in female flies chronically treated with rapamycin for five weeks display increase in climbing speed compared to untreated controls (Fig. 3e). Additionally, the time required to reach peak speed was reduced, suggesting improved agility and enhanced physical performance in rapamycin-treated female flies (Fig. 3f). Taken together, these findings are important for understanding how both intrinsic factors (ageing) and extrinsic factors (diet)contribute to physical decline over time. The age-related decline in performance mirrors sarcopenia and other muscle-related impairments seen in ageing mammals, while the high-sugar diet model highlights the impact of diet-induced metabolic dysfunction on physical ability. Furthermore, the ability of rapamycin to improve physical performance in aged flies shows that it can be used for the discovery and characterization of pharmacological interventions that prevent muscle decline. Together, these results suggest that *Drosophila* can serve as a valuable model to investigate the intersection of ageing, diet, and physical performance, providing insights into potential interventions for age-related decline and metabolic diseases.

### Exercise enhances endurance capacity and climbing performance

Regular endurance training is one of the most effective strategies to combat muscle loss, as it stimulates muscle growth and enhances strength. To investigate whether regular exercise can improve physical performance in *Drosophila*, male and female flies were subjected to a training protocol for five consecutive days. Males were exercised for 1 hour per day, while females were exercised for 30 minutes per day. Endurance capacity was then assessed using a climbing test over 3.5 hours, with climbing speed measured at the start and at 30-minute intervals. In females, exercised and non-exercised flies displayed similar climbing speeds during the initial phase of the endurance test (Fig. 4a,c). However, as the test progressed, exercised females maintained faster climbing speeds after the 1-hour time point and throughout the remainder of the 3.5-hour test, compared to non-exercised controls. Additionally, exercised females reached their top speed more quickly after one hour of endurance testing (Fig. 4e). These findings indicate that while exercise did not improve immediate climbing ability or the time to initial peak performance, it significantly enhanced endurance capacity, enabling flies to resist fatigue and sustain higher performance levels during prolonged physical exertion.

**Figure 4.**
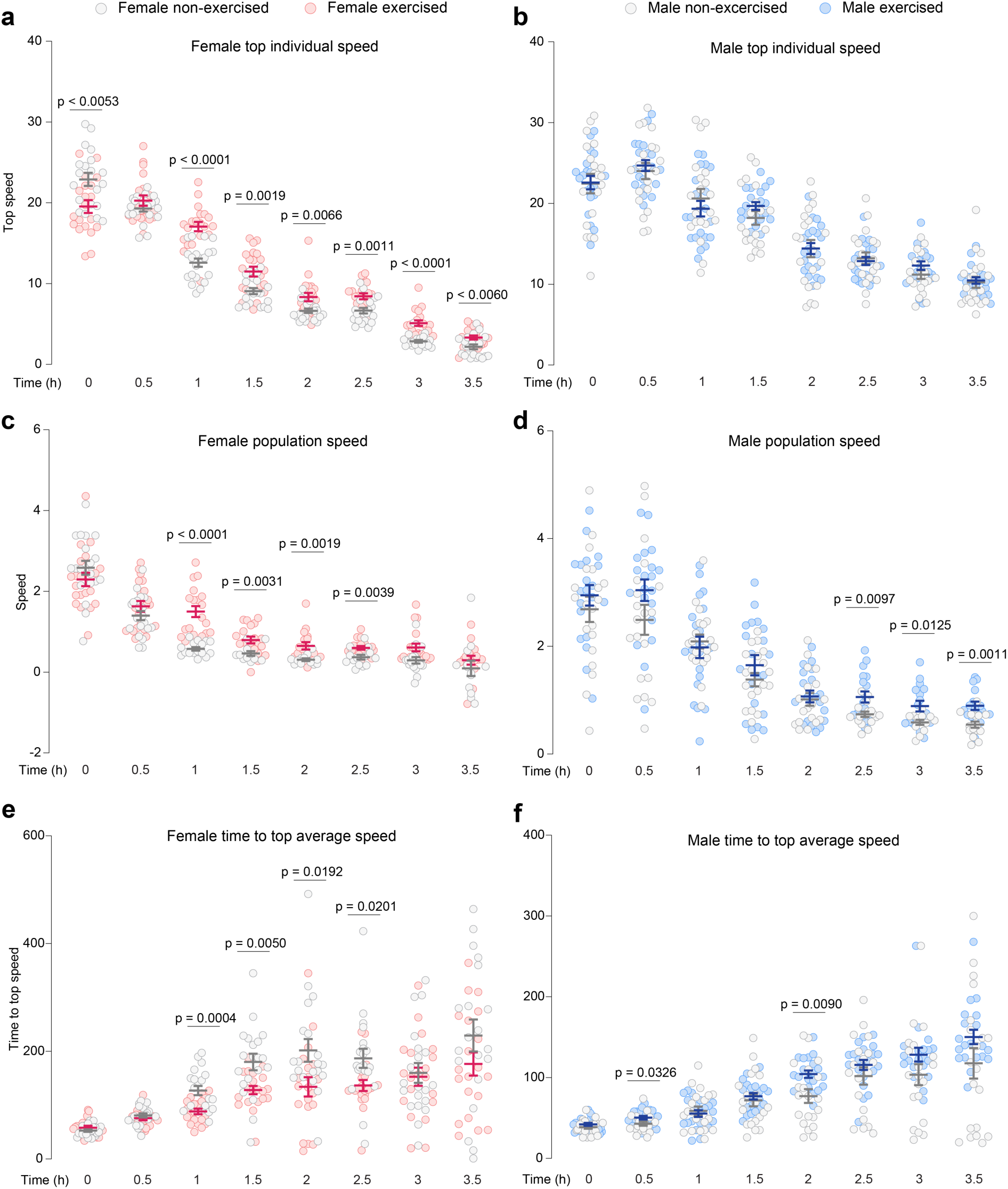
Effects of exercise on climbing endurance in male and female flies. Endurance capacity was measured in male and female flies following a five-day training protocol. Males were exercised for 1 hour per day, and females were exercised for 30 minutes per day. Climbing performance was assessed during a 3.5-hour endurance test, with performance recorded at the start and at 30-minute intervals. Top individual speed represents the highest climbing speed of individually tracked flies at each time point (a, b). Population speed reflects the maximum speed achieved by the group of flies at each time point (c, d). Time to top individual speed represents the time required for flies to reach their maximum climbing speed during each interval (e, f). Data are presented as individual data points with mean ± s.e.m. Statistical significance for pairwise comparisons between exercised and non-exercised flies was determined using two-tailed *t*-tests.

In males, the effects of exercise were more modest (Fig. 4b,d,f). Exercised males only displayed a significant benefit after 2.5 hours of endurance testing, achieving higher climbing speeds compared to non-exercised controls. This suggests that the current exercise protocol was insufficient to induce broader performance gains in males, or that males may require a different exercise regimen to achieve similar endurance improvements. Taken together, these findings demonstrate that endurance exercise enhances the ability to sustain climbing performance during prolonged physical activity, particularly in females, benefiting their ability to maintain performance over time rather than enhancing their immediate climbing capacity. Future studies should explore optimal exercise parameters for improving endurance in males and investigate the molecular mechanisms underlying sex-specific differences in exercise adaptations.

### Exercise promotes healthier activity patterns and induces a leaner body composition

We investigated the impact of endurance and chronic exercise on sleep and activity levels in male and female flies. Exercise protocols included a single day’s routine of endurance exercise for 3 hours or a chronic exercise regimen of one hour per day for males and 30 minutes per day for females for five days, with an additional 3-hour exercise session on the final day. Exercise led to significant reductions in both daytime and nighttime sleep in both sexes compared to non-exercised control animals (Fig. 5a– d). Female and male flies subjected to a single day of endurance exercise exhibited reduced sleep during both the following day and night compared to non-exercised controls, suggesting that a single bout of endurance exercise was sufficient to induce a pronounced reduction in sleep duration, similar to the effects of chronic exercise. These results demonstrate that both acute and chronic physical activity reduce sleep, indicating a systemic impact of exercise on rest behavior in *Drosophila*. Exercise also significantly increased daytime activity, as measured by beam-crossing events, in both male and female flies (Fig. 5e,f). In female flies, chronic exercise resulted in a notable increase in daytime activity, while a single day of exercise did not induce significant changes compared to controls (Fig. 5e). In contrast, both single-day and chronic exercise significantly increased daytime activity levels in males (Fig. 5f). Sleep and physical activity are well-established indicators of overall health and metabolic status in animals. The observed reductions in sleep and increases in activity patterns following exercise reflect physiological adaptations that may promote metabolic function and indicate potential health benefits.

**Figure 5.**
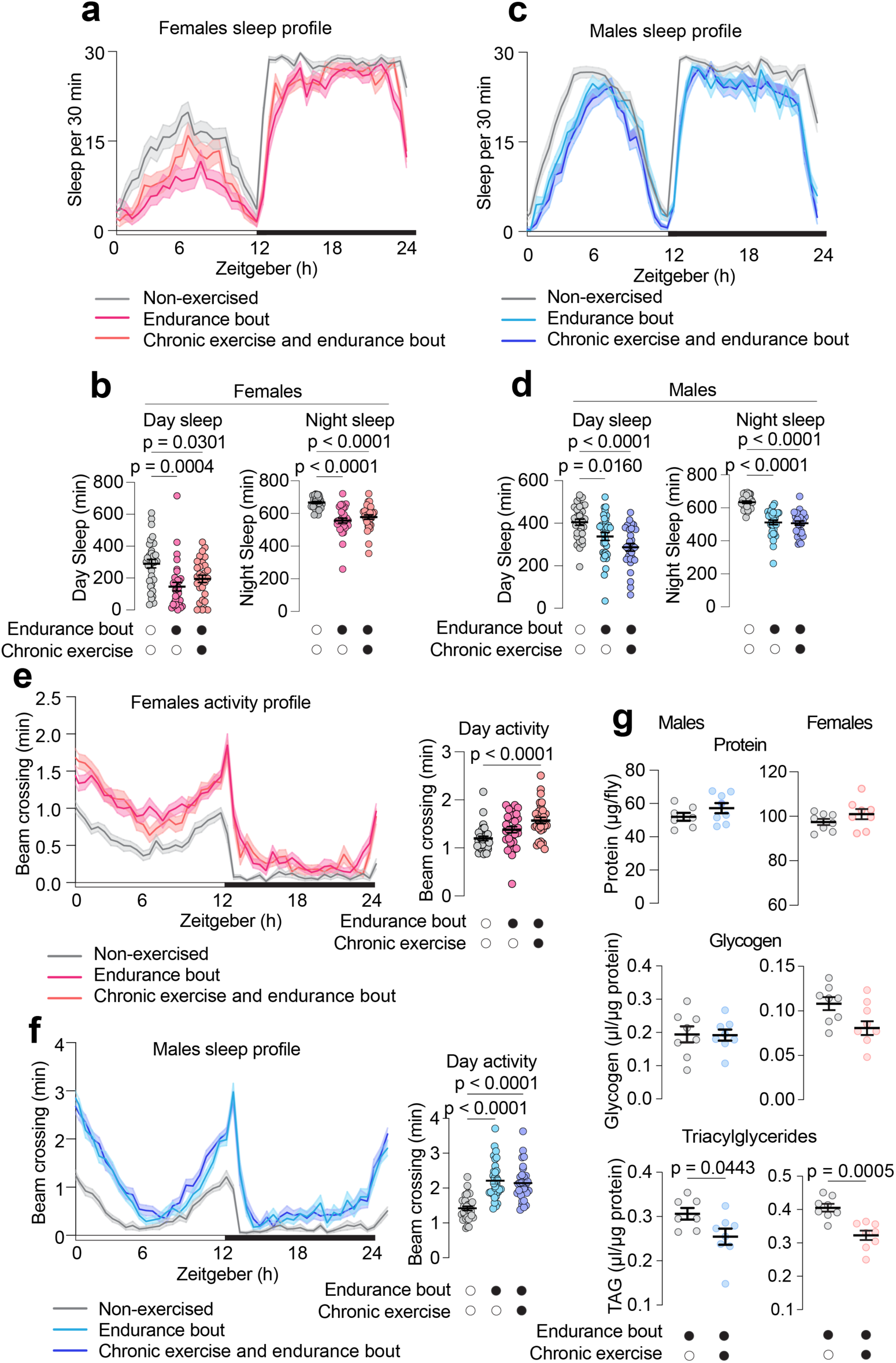
Effects of exercise on sleep, activity patterns, and body composition. Sleep and activity patterns were measured in non-exercised flies and animals subjected to either acute or chronic exercise protocols. The first condition consisted of a single 3-hour session, while chronic exercise included 1 hour per day for males or 30 minutes per day for females over five consecutive days, followed by a final 3-hour session. Twenty-four hours’ rest was allowed before these measurements. (a) Sleep over a 24-hour period in female flies across the three exercise conditions. (b) Daytime and nighttime sleep duration in female flies. (c) Sleep over a 24-hour period in male flies. (d) Daytime and nighttime sleep duration in male flies. (e) Daytime activity profiles measured as beam crossings per minute in female flies, with mean daytime activity levels shown in the bar plot on the right. (f) Daytime activity profiles in male flies, with mean daytime activity levels shown in the bar plot on the right. (g) Triacylglycerol (TAG), glycogen, and total protein levels in male and female flies subjected to chronic exercise compared to those that underwent a single day of exercise. Data are shown as mean ± s.e.m. Statistical significance was determined using one-way ANOVA with Tukey’s *post-hoc* test for panels (a–f) and two-tailed *t*-tests for panel (g).

To further investigate the long-term metabolic impact of chronic exercise, we measured the levels of triacylglycerol (TAG), glycogen, and total protein in male and female flies subjected to chronic exercise. The males were exercised for one hour per day for five consecutive days, while the females were exercised for 30 min per day for these days. On the sixth day both groups were subjected to an additional 3-hour endurance session and compared to flies that underwent a single day of endurance exercise (matching the last day of the “chronic exercise” animals). Our results indicate that both male and female flies subjected to chronic exercise exhibited significantly lower TAG levels, while the glycogen levels were similar (Fig. 5g). In addition, there were no significant changes in total protein levels (Fig. 5g). These findings indicate that chronic exercise promotes a leaner body composition in *Drosophila*, consistent with adaptations observed in mammalian models, in which exercise reduces fat stores while preserving lean body mass. The reduction in TAG levels suggests that chronic exercise induces long-term changes in energy storage and metabolism in flies, supporting the notion that physical activity enhances overall metabolic health across species.

### Exercise enhances both direct and insulin-dependent glucose uptake in *Drosophila* muscles

In mammals, exercise is known to increase glucose uptake into muscle, which supports the energy demands of physical activity. To investigate whether activity similarly enhances glucose uptake in *Drosophila* muscles, we measured the uptake of 2-NBDG (a fluorescent glucose analog) in male flies subjected to exercise. Flies were either left sedentary (non-exercised) or subjected to a 3-hour endurance-exercise session using the ClimbMaster apparatus. Following exercise, 2-NBDG was injected into the hemolymph (circulation), and the level of fluorescence in the musculature was measured 30 minutes post-injection by confocal microscopy. We separately assayed glucose uptake in the walking/jumping muscles and in other thoracic muscle tissues. Exercise significantly increased 2-NBDG uptake by the walking/jumping muscles by 36% compared to non-exercised flies (Fig. 6a). Similarly, glucose uptake in other thoracic muscles increased by 19% in exercised flies compared to non-exercised controls (Fig. 6b). These findings demonstrate that exercise enhances glucose uptake in *Drosophila* muscle tissues, similar to the well-documented effect observed in mammalian systems.

**Figure 6.**
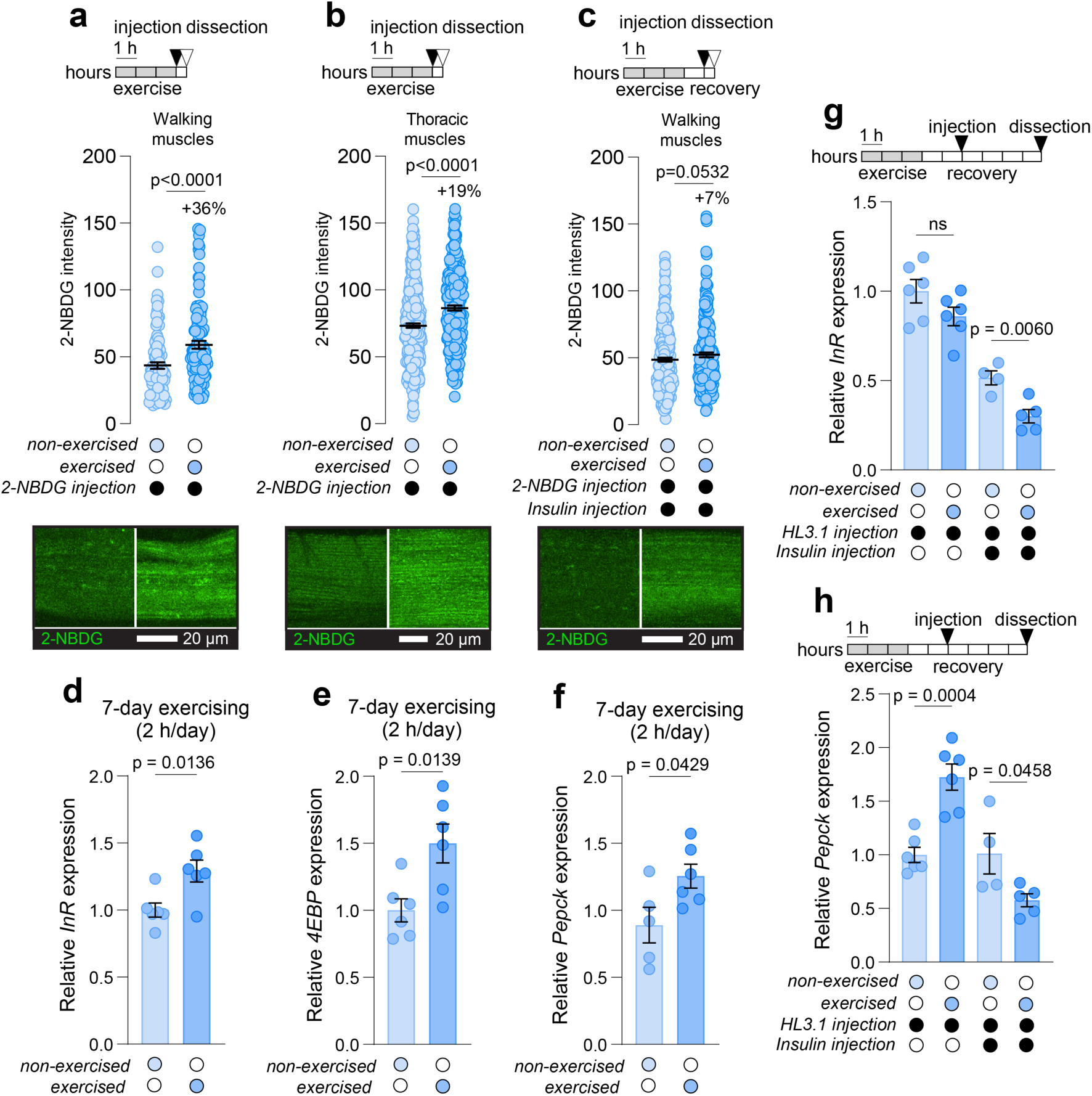
Exercise enhances glucose uptake and modulates insulin pathway gene expression in *Drosophila* muscle. Muscle glucose uptake was measured in male flies using 2-NBDG (a fluorescent glucose analog) injected into the circulation. (a) Glucose uptake in walking muscles of the thorax (b) Glucose uptake in other thoracic muscles. Flies were either subjected to a 3-hour endurance exercise session or kept under non-exercised control conditions. (c) Glucose uptake in walking muscles after a sedentary period or a 3-hour exercise session followed by a 1-hour rest period, with insulin injected together with 2-NBDG. Representative images of 2-NBDG fluorescence in muscle fibers are shown below each panel. Flies in all groups were kept on starvation medium (1% agar) for 3 hours prior to injection, and muscles were dissected 30 minutes post-injection. Data in panel (c) were log-transformed for statistical analysis. (d–f) Transcript levels of *Insulin receptor* (*InR*) (d), *Eukaryotic translation initiation factor 4E-binding protein* (*4EBP*) (e), and *Phosphoenolpyruvate carboxykinase* (*Pepck*) (f) in thoracic muscle from flies subjected to chronic exercise (7 days, 2 hours per day) or no exercise. Gene expression was measured by qPCR from dissected muscle tissue from males. (g, h) Expression of *InR* (g) and *Pepck* (h) in muscle from flies subjected to acute exercise (3 hours), followed by 1 hour of recovery and injection with either saline (HL3.1) or insulin in saline. Gene expression was measured 4 hours post-injection. Data are presented as mean ± s.e.m. Two-tailed t-tests were used for pairwise comparisons. ns indicates non-significant differences. Data are shown as mean ± s.e.m. Statistical significance was determined using two-tailed *t*-tests for pairwise comparisons.

To further investigate whether exercise enhances insulin-dependent glucose uptake, which is typically elevated within one hour of exercise in mammalian models, we subjected male flies to a 3-hour exercise session followed by a 1-hour rest period. After the rest period, insulin was injected together with 2-NBDG. Our results indicate that exercise enhanced insulin-mediated glucose uptake in *Drosophila* leg muscles (Fig. 6c, *p* = 0.0532). These findings suggest that, as in mammals, exercise primes *Drosophila* muscles to become more responsive to insulin. Taken together, these results indicate that *Drosophila* muscles respond to exercise by increasing glucose uptake both directly and in an insulin-dependent manner, highlighting the conserved nature of exercise-induced glucose regulation across species.

### Exercise reduces insulin pathway activity and enhances insulin responses in muscle

One of the key effects of physical activity in mammals is its lowering of insulin levels and improvement of insulin sensitivity, which facilitates glucose uptake into muscle tissue and helps regulate blood glucose levels^25^. To examine whether exercise induces similar effects in *Drosophila*, we first subjected male flies to chronic exercise (7 days, 2 hours per day) using the ClimbMaster apparatus and measured the expression of *Insulin receptor* (*InR*) and *Eukaryotic translation initiation factor 4E-binding protein* (*4EBP*), which are direct FOXO targets and are thus repressed by insulin signaling^26,27^. We observed that chronic exercise significantly increased the transcript levels of *InR* and *4EBP* in the muscles, indicating reduced basal insulin pathway activity, consistent with lowered insulin levels in response to long-term exercise (Fig. 6d,e). We also assessed effects on metabolism by measuring the expression of *Phosphoenolpyruvate carboxykinase* (*Pepck*), a key enzyme in gluconeogenesis, which is typically repressed by insulin to reduce glucose production^28^. *Pepck* transcription was significantly elevated in chronically exercised flies, consistent with reduced insulin signaling and an increased demand for glucose production during prolonged physical activity (Fig. 6f). These results suggest that chronic exercise leads to a sustained increase in the expression of insulin-repressed genes in muscle, reflecting adaptive changes in energy metabolism that support repeated bouts of physical activity.

Insulin sensitivity in humans is known to increase during the early post-exercise period, typically peaking within 1-3 hours after exercise, and can remain elevated for hours^29^. To capture a *Drosophila* post-exercise window comparable to those in human studies, male flies were subjected to a single 3-hour endurance exercise session, followed by 1 hour of recovery before injection with insulin or saline. Importantly, flies were kept without access to food during both the exercise and the 1-hour recovery period to reduce endogenous insulin release, mimicking the fasting conditions commonly used in human post-exercise insulin sensitivity studies. Non-exercised control flies were similarly kept without food for 4 hours to ensure consistent conditions across all groups. We found that *InR* expression was more strongly repressed by insulin in muscles from exercised flies compared to non-exercised controls, indicating an enhanced muscle insulin responsiveness after exercise (Fig. 6g). We also assessed the expression of *Pepck* and found that in saline-injected flies, *Pepck* expression increased following exercise, consistent with the ability of physical activity to reduce insulin signaling and increase the demand for glucose production (Fig. 6h).

### Exercise modulates insulin signaling and protein phosphorylation patterns in *Drosophila* muscle

To further investigate insulin signaling after exercise, we performed a Western blot to measure the level of phosphorylated AKT (pAKT), an early marker of insulin signaling activation, one hour after a 3-hour exercise regimen, as mammalian muscle typically exhibits increased insulin-stimulated glucose uptake during the hours following exercise. Both exercised and non-exercised animals were placed on starvation medium during the protocol to prevent feeding-induced insulin release. Following the exercise session, animals were divided into two groups: one receiving a mock injection and the other injected with human insulin. Animals that received a mock injection displayed minimal pAKT signal in both the non-exercised and exercised conditions. In contrast, animals injected with human insulin exhibited a strong pAKT signal in both conditions, indicating effective insulin pathway responses at the level of AKT, which is several steps downstream of the insulin receptor in the signaling cascade (Fig. 7a). Although a non-significant trend towards somewhat increased pAKT abundance after exercise was observed, this similarity in response suggests that exercise might affect mechanisms downstream of AKT in the insulin pathway or sugar-uptake mechanism, as it does in mammals, in which exercise is believed to enhance insulin-dependent glucose uptake in part via phosphorylation of TBC1D4 (also known as AS160), a Rab-GTPase-activating protein involved in GLUT4 translocation^30^.

**Figure 7.**
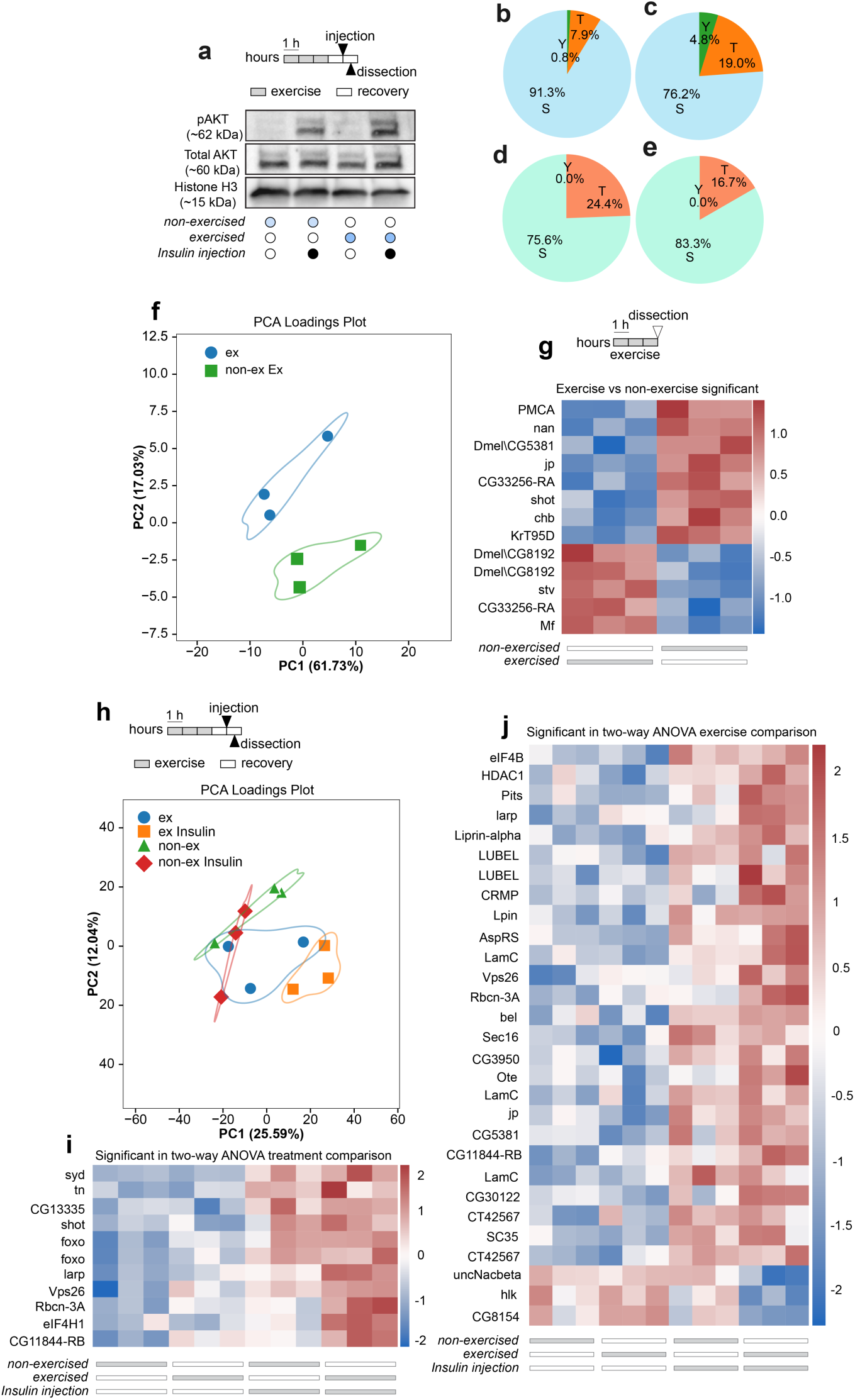
Phosphoprotein analyses of the impact of insulin and exercise. (a) Western blot analysis showing activation of the insulin pathway through phosphorylation of Akt in the thoraxes of male flies subjected to 3 hours of endurance exercise or kept under non-exercised control conditions. Following a 1-hour rest period, animals were injected with insulin or mock solution. An additional 30-minute rest period was allowed to facilitate the insulin response before dissection. (b) Composition of phosphorylated residues identified in all phosphoproteomic analyses. (c) Composition of phosphorylated target sites altered between exercised and non-exercised, never-injected animals. (d) Similar to (b), but representing changes induced by exercise in injected animals, regardless of injection content. (e) Similar to (d), but representing changes between insulin and mock (HL3.1) injections, regardless of exercise status. (f) Principal component analysis (PCA) of exercised and non-exercised animals that were not injected. Outlined boundaries represent 50% probability contours calculated via kernel density estimation. (g) Proteins exhibiting significantly different modification profiles between exercised and rested animals (related to panel f). (h) PCA of animals that were either exercised or rested and then injected with either HL3.1 or HL3.1+insulin. (i) Proteins exhibiting significantly different modification profiles when comparing insulin injection to mock injection, regardless of exercise status. (j) Similar to panel (i), but comparing exercised and rested animals, regardless of injection status. Data are presented as mean ± s.e.m. Two-tailed t-tests were used for pairwise comparisons. ns indicates non-significant differences.

To perform deeper analyses of exercise-induced biochemical changes, we conducted quantitative phosphoproteomic studies to examine the impact of exercise and insulin on post-translational modifications in *Drosophila* muscle tissues. To assess the immediate effect of exercise on muscle, phosphoproteomic profiles of animals subjected to 3 hours of endurance exercise were compared to those of rested controls. In a second experiment, examining post-exercise muscle phosphosite responses and whether insulin responsiveness is modulated after exercise, we examined phosphoproteomic profiles one hour following the 3-hour exercise regimen in animals that had been injected with insulin or vehicle. Among all catalogued phosphosites, serine residues accounted for the majority (>90%), with threonine (∼8%) and tyrosine (<1%) representing a smaller fraction (Fig. 7b-e). Notably, phosphosites differing between exercised and non-exercised animals exhibited an increased proportion of threonine and tyrosine residues, implicating potential roles for tyrosine and threonine kinases in exercise-induced signaling.

Principal-component analysis (PCA) revealed that exercised and non-exercised animals were clearly separable based on their phosphoproteomes, indicating that endurance exercise induces coordinated changes in phosphorylation patterns in muscle tissue (Fig. 7f). Among the proteins significantly altered by exercise, Mitofusin (Mf; homolog of MFN2), which regulates mitochondrial fusion and is linked to insulin sensitivity^31^, exhibited phosphorylation changes in response to exercise (Fig. 7g). These modifications likely reflect enhanced mitochondrial function, a hallmark of endurance training in both flies and mammals.

PCA of data from the second experiment, involving insulin injection, indicated that exercised animals responded distinctly to insulin along the PC1 axis compared to rested controls (Fig. 7h), suggesting that exercise modulates muscle pathway responses to insulin. Furthermore, selected phosphoproteins significantly altered by insulin stimulation (Fig. 7i) included FOXO, a key downstream component in the insulin signaling pathway; eIF4B, a protein indirectly regulated by FOXO (via its transcriptional effects on *eIF4B-BP1*); and Vps26, a protein involved in endosomal sorting and thus potentially affecting GLUT4 translocation. Indeed, loss of Vps26 has been observed to decrease GLUT4 levels, suggesting that Vps26 plays a role in maintaining GLUT4 expression and, consequently, insulin-dependent glucose uptake^32^. In post-exercised muscles, eIF4B exhibited altered phosphorylation, further implicating exercise as a modulator of insulin signaling and protein synthesis, as did Sec16, which plays a role in regulating GLUT4 trafficking and thereby affects insulin-stimulated glucose uptake in mammalian models^33^ (Fig. 7j). Closer analysis also revealed that exercise potentiates insulin-stimulated phosphorylation of eIF4B (*p* = 0.012) and Vps26 (*p* = 0.030). These findings indicate that exercise modulates phosphorylation patterns in pathways responsive to insulin and involved in glucose uptake in *Drosophila* tissues.

## Discussion

The ClimbMaster, paired with a custom automated video-analysis pipeline, enables precise and reproducible studies of the physiological and molecular adaptations to exercise in *Drosophila*. By integrating automated hardware with advanced software, the apparatus ensures consistent operation with precise timing, facilitating high-throughput experiments with minimal variability. The inclusion of a MATLAB-based video analysis script further enhances the system’s utility, providing detailed performance metrics such as climbing speed, endurance, and consistency. Together, these tools offer a robust and scalable platform for exploring the effects of exercise under diverse genetic, dietary, and environmental conditions, making it a versatile system with virtually unlimited potential for scaling.

The ClimbMaster provides a high-throughput platform that can be used to screen thousands of drug candidates in a matter of weeks or months to identify anabolic compounds that improve muscle function and performance or enhance the biological responses to exercise. This system enables rapid functional assessment of genetic and molecular pathways identified in human studies that are regulated by physical activity^34^. For example, exercise-induced phosphorylation sites or redox-regulated cysteines that change with physical activity and aging^34,35^ can be genetically manipulated in flies using CRISPR-based technologies^36^ and paired with the high-throughput assessment of the ClimbMaster to evaluate their impact on physical performance, muscle physiology, insulin sensitivity, and glucose uptake. Such manipulations can be easily achieved at a large scale by genetically engineering fly genes or by humanizing the fly through replacing endogenous fly genes with modified versions of human genes^37–39^. The system also allows for the testing of human genetic variants that are associated with metabolic disorders, physical activity, muscle function, and sarcopenia. There is currently a significant unmet need for the functional characterization of human variants identified through genomic studies, which cannot be addressed by existing *in-vitro* or *in-vivo* models. The ClimbMaster offers unprecedented functional assessment capabilities *in vivo*, enabling large-scale genetic and pharmacological screens that can drive progress in this field. Our results show that the system recapitulates key aspects of mammalian exercise physiology, including the impact of physical activity on insulin sensitivity and glucose metabolism, further supporting its translational relevance.

Exercise enhances insulin-dependent glucose uptake in mammalian and human muscles^40,41^. Since we demonstrate that key features of the exercise-induced increase in muscle insulin sensitivity and insulin-mediated glucose uptake are conserved between flies and mammals, the ClimbMaster can also be used to characterize the mechanisms underlying insulin resistance based on human studies, such as those identifying changes in skeletal muscle linked to insulin action^42^. Insulin resistance, a key driver of T2D, occurs when muscle, fat, and liver tissues become less responsive to insulin, causing the pancreas to compensate by producing more insulin^9^. Over time, this compensatory mechanism fails, leading to chronically elevated blood glucose levels and T2D. Insulin resistance is characterized by impaired glucose uptake by muscle tissues, leading to elevated blood glucose levels and subsequent metabolic dysregulation. By permitting measurement of muscle glucose uptake and insulin sensitivity in flies exposed to varying diet-and-exercise protocols and diverse genetic manipulations, this system offers a powerful tool to decipher how specific pathways and genetic variants identified in personalized human analyses^42^ contribute to insulin resistance. Understanding these mechanisms is essential for developing effective interventions to combat T2D and other metabolic disorders. This capability highlights the broad applicability of the system for studying metabolic health, alongside its utility in muscle function and physical performance research.

Beyond exercise and muscle function, the ClimbMaster has broad applicability in other disciplines, including the study of neurodegenerative diseases that impair physical abilities. Diseases such as Parkinson’s disease, amyotrophic lateral sclerosis (ALS), and Huntington’s disease are characterized by declining motor function, and the ClimbMaster can be used to assess physical performance and health in these models^43^. As with muscle function, the system can be used to conduct large-scale genetic screens and drug discovery efforts to identify interventions that improve mobility and overall health in neurodegenerative-disease models.

Sleep and activity patterns are widely recognized as key indicators of healthspan and aging^44,45^, which refer to the period of life spent in good health, free from chronic diseases and disabilities. We show that exercise reduces sleep and increases activity, indicating improved behavioral outputs linked to healthspan and aging. Combining the ClimbMaster exercise platform with other high-throughput systems that allow monitoring of sleep, activity, feeding, and other behaviors [such as the *Drosophila* Activity Monitor (DAM) and FLIC systems^46^], provides a unique opportunity to gain insights into the effects of exercise on healthspan and aging. These approaches are only feasible in *Drosophila* and enable the simultaneous assessment of physical performance, sleep, and activity patterns, which are critical indicators of overall health, aging, and longevity. While physical performance measurements from the ClimbMaster provide direct insights into muscle function and endurance, sleep and activity metrics offer a more comprehensive picture of healthspan and aging effects that are only possible through *in vivo* studies. Thus, the ClimbMaster provides a powerful system bridging the fields of exercise and aging, offering valuable insights into how physical activity influences aging-related processes in humans.

In summary, the ClimbMaster’s ability to induce exercise and to monitor behavioral outputs, paired with the molecular-genetic approaches available in *Drosophila*, makes it a powerful tool for investigating the genetic and environmental factors that influence muscle health, metabolic function, and physical performance. Its scalability and precision enable detailed studies that are infeasible or impractical in mammalian models, facilitating high-throughput screens to uncover novel therapeutic targets. By providing a comprehensive platform for studying the effects of exercise and related interventions, the ClimbMaster holds the potential to accelerate discoveries in exercise physiology, metabolic health, aging research, and disease biology and inform the development of therapeutic strategies for sarcopenia, metabolic disorders, and central nervous system disorders, contributing to the broader goal of improving healthspan and quality of life across the lifespan.

## Methods

### *Drosophila* rearing and maintenance

Flies were reared on a standard cornmeal-based diet containing 82 g/L cornmeal, 60 g/L sucrose, 34 g/L yeast, 8 g/L agar, 4.8 mL/L propionic acid, and 1.6 g/L methyl-4-hydroxybenzoate. Except for the experiments of Fig. 1, which will be described in detail elsewhere, all experiments were conducted using *w^1118^* flies, the genetic background for most transgenes, which were maintained at 25 °C with 60% relative humidity under a 12-hour light/dark cycle. Upon eclosion, flies were transferred to an adult-specific cornmeal-free diet consisting of 90 g/L sucrose, 80 g/L yeast, 10 g/L agar, 5 mL/L propionic acid, and 15 mL/L of 10% methyl-4-hydroxybenzoate in ethanol. This diet was provided 4–6 days before experiments, and flies were maintained on it until testing. Adult mated females and males, aged 5–10 days, were used for all experiments except for the assessment of physical performance during aging in which the age of the animals is written in the figure. To ensure optimal conditions, flies were provided with fresh food every two days throughout the experimental period. For the experiments with high sugar diet, the adult specific food was modified to contain 20% sugar. The rapamycin (Termo Scientific #15434849) concentration used was 10 µg/ml dissolved in dimethyl sulfoxide and it was mixed in the adult-specific food during the cooking process after cooling down to bellow 40°C to avoid degradation.

### Exercise and climbing analysis protocol

For experiments, flies were transferred to fresh standard fly vials, each containing approximately 20 animals, and tightly and securely fixed in the ClimbMaster grid. The exercise protocol typically consisted of 1 to 2 hours of repeated climbing per session, with automated tap-down movements occurring at 8-second intervals. These controlled displacements triggered the flies’ innate negative geotaxis behavior, effectively simulating endurance exercise. To assess endurance, flies were subjected to extended exercise periods of up to 4–5 hours, maintaining the 8-second tap-down interval. Performance was monitored at 30-minute intervals by recording 10 consecutive rounds of climbing, each followed by a 20-second interval to allow the flies additional climbing time. Recorded videos were analyzed using custom-written software to quantify climbing speed, endurance, and consistency across experimental groups. The software, designed for reproducibility and efficiency, generates detailed metrics and will be provided upon request.

### Automated video analysis and climbing performance quantification

To efficiently and reproducibly quantify climbing performance in *Drosophila*, we developed a custom MATLAB script tailored to analyze video recordings of the ClimbMaster assays. This automated approach enables high-throughput processing of repeated climbing tests, providing detailed and accurate performance metrics. The algorithm proceeds as follows.

1. **Metadata setup:** the user selects one or more video files of any common format (QuickTime MOV, MP4, AVI).
2. If enough memory is available, the entire video is loaded at once; otherwise, each frame is loaded and analyzed separately. Therefore, videos may be of any size or length.
3. The user is presented with the first frame of the video as a reminder of its contents.
4. Initial setup data is obtained from the user, primarily including the number of vials that are present.
5. The names/genotypes of each vial, and the number of flies in each, is input.
6. A region of interest is drawn across the video frame to enclose the region to be monitored, divided into containers for each vial.
7. As a means to normalize for different levels of zoom or video resolutions, the user sets a scale bar to some object of fixed size that is used for every video, such as the width of a vial or a piece of the machine. All measurements of distance or speed that are output in the analysis are scaled to this standard.
8. **Identifying test intervals:** with these initial parameters set, the program scans the entire video and quantifies the amount of change occurring from one frame to the next.
9. Frames containing extremely large amounts of motion are interpreted as those in which the apparatus itself is moving to “bang” the flies to the bottom of the vial. The user is requested to set a triggering value for motion that indicate these slams (or in batch mode, the value is selected automatically). A frame at which the measured amount of motion drops below this line marks the beginning of a test interval, and the time at which the motion exceeds this limit again marks the end of that interval.
10. Settings for fly identification are requested from the user. These values indicate the maximum and minimum sizes of objects that will be deemed “flies”, as well as aspects of their shape. These values are scaled using the factor specified at step #7, so these values do not need to be adjusted often.
11. In the steps below, the segmented vial compartments within the user-selected region of interest are processed separately.
12. A new background image, from which the flies have been mathematically removed, will be created for each interval. Making the assumption that flies are dark, and the background is light, frames dispersed across the interval are combined using a maximum-brightness projection.
13. The fly-free background is removed from each frame within the interval, leaving the flies as detectable objects.
14. Remaining objects, including flies but perhaps other things, are segmented out of the image, and their size and other metrics are determined. Objects that meet the parameters set in step 10 are deemed to be flies, whereas other objects are discarded. If the user has requested that videos or images be output during processing, the objects will be color-coded as flies or to indicate which parameter led to their being discarded.
15. As a measure of group fly position that does not require knowledge or detection of the number, identity, or motion of the flies, the center of mass of the collection of FLOs (that is, weighted by their size) is determined in each frame.
16. The position, size, and shape of each fly will be recorded for later track reconstruction.
17. Once all the flies have been classified in every vial, in each frame of each interval, the data will be saved so that it can be reprocessed later if desired.
18. Tracking fly motion: The user will be asked for parameters by which to adjust the tracking algorithm. These values are also scaled according to the measure in step 7 as appropriate, so they generally do not require adjustment. (In batch mode, this query will be made immediately after the first parameter input, rather than at this point.)
19. If the user’s video-recording frame-rate is adequate (*e.g.*, 25 or 30 frames per second), it is very likely that for each fly “A” in video frame *N*, the fly in frame *N-1* that is closest to the position of fly “A” is indeed still fly “A”.
20. Fly “A” in the last frame of the interval is traced backwards through the recorded position data until its track ends (or rather, begins). As the track is reconstructed, this fly is removed from the data to prevent double-tracking. The track is plotted into an output graphic, with a circular marker placed at the latest position in each track. The size of this marker is proportional to the amount of time spent at that position – large markers indicate that the FLO reached its position early in the interval and had not moved in a relatively long time.
21. Remaining flies in the last frame of the interval are likewise traced back through the data until no flies remain in this frame.
22. The analysis begins again at each preceding frame in the interval, if any flies are present in that frame (that is, if they have not been removed during path tracing)., until no untracked flies remain in the data set.
23. The frame-to-frame vertical motion of all the flies in vial is averaged for every frame in the interval.
24. A variety of data are output for each vial, either for all intervals separately or as an average across intervals, discussed further below.
25. After the tracking has completed, a one-phase exponential decay function is fit to the “center of mass over time” curve that reflects the population performance:
26. y = (Y0–Plateau)e^-Kx^ + Plateau
27. The first derivative of this function gives the slope of the curve at each point – that is, the rate at which the center of fly mass is moving at any time, or to put it most succinctly, the climbing speed of the whole population.
28. dy/dx =-K(Y0–Plateau)e^-Kx^
29. The slope is by default calculated at two time points: (1) at 30 frames after the beginning of the interval, an empirically chosen time; and (2) at the time when the population has climbed halfway up from its minimum height to its maximum height. Because the function is an exponential decay, this time point is mathematically meaningful and represents one “half-life time constant.”
30. If the program is not in batch mode, the user is asked whether any of the default analyses need adjustment. If so, the curve-fitting parameters can be adjusted interactively.

### Data outputs

The most useful numerical data are output into a comma-separated-value (CSV) file, “Scaled Coefficients,” in a folder named after the selected video. Within this file, the following values are given for each vial, during each interval. The most salient are bolded. *Fit coefficient 1 (Y0)* – the starting height of the population. *Fit coefficient 2 (Plateau)* – the height that the population will reach in infinite time. ***Fit coefficient 3 (decay constant K above)*** – the parameter of the exponential function; for two tests in which the differences Plateau-Y0 are the same, a larger decay constant indicates faster vertical “center of mass” motion (assuming the distance between Y0 and Plateau are equal). *Half life* – the time required for the population average to climb halfway from the minimum position to the maximum height; mathematically, the reciprocal of the decay constant. *Fit R-squared* – a measurement of the match between the data and the fit curve. ***Total activity*** – a measure of the amount that the fly images change from frame to frame. Broadly interpretable as a measurement of motion in the vial. ***Slope at point X*** – the slope of the fit curve at frame 30 (or whichever frame is specified by the user) – the climbing rate of the “center of fly mass”. *X position for slope calculation* – the frame at which the slope is calculated (30, unless this is changed). *Y position at slope calculation* – the population height at this time. *X position at Y midpoint* – the frame at which the population height reaches the halfway point between Y0 and Plateau; one “half-life”. *Y position at Y midpoint* – the population height here, halfway between Y0 and Plateau. ***Slope at this midpoint*** – the climbing speed of the population at this midpoint time. *95th percentile climbing speed* – the 95^th^ percentile (that is, higher than 95% of values) of climbing speed (where “climbing speed” is the average of individual fly movements from frame N to frame N+1). Note that this figure is affected by the length of the observation, whereas the following measurement is not. ***Max of moving-average climbing speed*** – the peak of the averaged individual speeds derived from tracking. Two related measurements are output: the speed averaged across the number of flies detected in the vial, and the value averaged across the number of flies that the user input. ***Time to peak climbing*** – the video frame within the interval at which this top speed was reached.

#### Image outputs

PNG and PDF images will be output.

*All Pixel Motion* – each point represents the amount of frame-to-frame change in one of the vials, in one of the intervals. Points are color-coded by vial but not by interval. *All Tracked Climbing* -- each point represents the amount of frame-to-frame change in one of the vials, in one of the intervals. Points are color-coded by vial but not by interval. A horizontal line indicates the 95^th^ percentile values reported in the CSV file. *All Tracks* – the reconstructed tracks of individual flies are shown, with an track-end marker representing the amount of time the fly had remained still at that position when the track ended. *Average Pixel Motion* – lines color-coded per vial represent the average between the points in the “All pixel motion” file. *Average Tracked Climbing speed* – color-coded lines indicate the climbing speed of tracked fly-like objects, averaged across the detected FLOs and then across all experimental intervals. *Center of fly mass, all intervals* – the vertical position of the population, with each interval shown separately and each vial color coded. *Center of fly mass, averaged across intervals* – similar to the image above, but with a single line per vial indicating the center at each point, averaged across the several test intervals. *Cumulative pixel motion* – “Average pixel motion” cumulated over time. *Cumulative tracking* – “Average tracked climbing speed” integrated over time – interpretable as the position of all the individual flies, averaged over time; as such, it provides information similar to that given by “Center of fly mass, averaged across intervals”, but it is calculated independently. *Fit for vial X, interval N* – information about the decay function that was fit to the climbing profile in Vial #x for the performance in the Nth interval. *ROI* indicates the selected region that was analyzed during the program run. *Running-Average All Climbing* – The data shown in “All tracked climbing” but processed with a running average to smooth frame-to-frame variation.

**CSV files of numerical data** are output for analysis in other software such as Microsoft Excel or GraphPad Prism. *Center of fly mass* – CSV file cognate with “Center of fly mass, all intervals” image. *ClimbingAverages* – CSV file cognate with “Average tracked climbing speed” image. *Pixel action list* -- CSV file listing the amount of detected video change within each vial, within each frame of each interval. This is cognate with the “All pixel motion” image.

#### If the user elects to output interim processing videos

*MotionVideo.mp4* indicates the frame-to-frame pixel changes quantified as “pixel motion” in the data above. *DetectionVideo.mp4* shows all the detected objects, color-coded by their processing fate: fly-like objects are green, and other segmented features are colored to indicate why they are deemed “not fly-like” – too big (purple), too small (red), too long (white), too skinny (yellow). A horizontal blue line indicates the center of mass of the green pixels. *TrackingVideo.mp4* indicates the tracked motion of the green FLOs. Each tracked FLO is given a purple vector enlarging its movement from the prior frame to the present frame. The white bar indicator at the center of each vial indicates the climbing motion at that time, averaged across all detected FLOs. FLOs outside the tracking window within each vial are colored a darker green; these animals are not analyzed in the “tracking” data but do contribute to the “center of mass” data.

#### Statistical comparisons

The script performs ANOVA evaluations between the vials with regard to 95^th^-percentile climbing speed, decay constants, half-life, pixel motion, population-height slopes at the user-selected frame, and population-height slopes at the midpoint between Y0 and Plateau. These are output as “Chart of…” Further analysis is left as an exercise for the user.

#### Internal files

*Data0.mat* is a metadata file containing vial names and ROI data. *Data1.mat* contains the motion data used to select intervals. *Data2.mat* contains the data on detected fly-like objects. *Data3.mat* contains fly-tracking data. *Data4.mat* contains derived performance data. *Dataset.mat* contains further data that can be aggregated using the combiner script.

#### Combiner script

Often we analyzed experiments that contained multiple vials of the same genotype, and we often performed several experiments that we wished to pool and compare with one another. The Combiner script performs these functions. After all relevant videos are analyzed, the Combiner script loads their associated “Dataset.mat” files to retrieve the respective vial names and performance data. The user is prompted to name the groups of data that the script will create (e.g., “all the vials containing males”), and then the vials to be included in each group are selected from a list of all the vials across all the videos. The performance data are regrouped and compared through ANOVA. Relevant charts are output into the MatLab current directory for simple inspection, along with a CSV file of numerical data, which can be transferred to a columnar dataset in GraphPad Prism, for example, by a “paste with transpose” operation, or inserted into a “grouped” (two-way ANOVA) spreadsheet by a simple paste.

### Sleep and activity assays

Sleep and activity patterns were measured using the *Drosophila* Activity Monitor (DAM) system (TriKinetics, Waltham, MA). Individual male or female flies (7–10 days old) were placed into 2-mm inner-diameter glass tubes mounted in DAM2 monitors. One end of each tube was sealed with a detachable 250-μL PCR tube containing 90 μL of feeding medium, while the other end was plugged with plastic foam to secure the flies. The feeding medium consisted of 5% sucrose in 1% agar dissolved in water to provide hydration and a carbohydrate source during the monitoring period. Monitors were housed in an incubator set to 25 °C with 60% humidity on a 12-hour light/dark cycle. Data from the first day and first night immediately following exercise were discarded to account for post-exercise recovery. Data from the following day and night period were analyzed to assess the effects of exercise on sleep and activity. Sleep was defined as five consecutive minutes of inactivity, as detected by infrared beam crossings in the DAM system, as is customary in the field. Activity and sleep data were recorded continuously for one 24-hour period. Beam-crossing events were used to quantify activity levels, and sleep duration was calculated as the total time spent inactive during both daytime (lights on) and nighttime (lights off) phases of the light/dark cycle. Data were analyzed using custom MATLAB scripts (The MathWorks, Natick, MA).

### 2-NBDG uptake assay in *Drosophila* muscles

Glucose uptake by *Drosophila* muscle tissue was measured using 2-NBDG (2-[N-(7-nitrobenz-2-oxa-1,3-diazol-4-yl) amino]-2-deoxyglucose), a fluorescent glucose analog that can be transported into cells but only metabolized in limited ways afterwards. Male flies were transferred to starvation medium (1% agar in water) for 3 hours prior to injection to prevent food intake and associated endogenous insulin release. Experiments in Figure 6a,b: For the exercise condition, flies were subjected to a 3-hour endurance exercise session using the ClimbMaster apparatus, while non-exercised control flies remained on starvation medium during the same period. Following exercise or the control period, each fly received a thoracic injection of 100 nL (roughly 10% of the endogenous fluid content) of either 5-mM 2-NBDG in PBS or PBS alone. The injection solution was prepared from a 50-mM stock solution of 2-NBDG in ethanol (100%), which was diluted in PBS to achieve a final concentration of 5 mM. This resulted in an estimated final concentration of 0.5 mM 2-NBDG in the hemolymph. The ethanol concentration in the injection solution was 10%, resulting in 1% ethanol in the hemolymph. Flies were kept on starvation medium during the 30-minute glucose uptake period following injection. Figure 6c: insulin-induced glucose uptake was assessed by subjecting flies to a 3-hour exercise session on starvation medium, followed by a 1-hour recovery period on the same medium. After the recovery period, flies were injected with 100 nL of a solution containing 5 mM 2-NBDG and 100 µg/mL human insulin (Sigma-Aldrich, I2643) in PBS. This resulted in an estimated final concentration of 0.5 mM 2-NBDG and approximately 10 µg/mL insulin in the hemolymph. Muscle tissue was dissected 30 minutes post-injection, including thoraxes and legs, with wings, gut, and ventral nerve cord (VNC) removed to isolate the muscles. In all experiments, dissections were performed in ice-cold PBS to preserve tissue integrity. Muscle samples from the side of the body contralateral to the injection were fixed in 4% paraformaldehyde (PFA) for 25 minutes at room temperature, rinsed with PBS, and washed three times for 5 minutes each. Samples were mounted in ProLong Gold Antifade Mountant (Thermo Fisher) without additional nuclear staining. Glucose uptake was assessed by visualizing 2-NBDG fluorescence in intact tissue using confocal fluorescence microscopy (Zeiss LSM-900, 20x objective, 2-micron Z spacing). Care was taken to select intact muscle fibers to avoid artifacts from broken fibers or glucose leakage. Separate measurements were taken for walking/jumping muscles and other thoracic muscles. In ImageJ, Z-stacks were max-projected, and the average fluorescence intensity after background subtraction was recorded at representative locations within each sample.

### Transcript analysis using qPCR

Muscle tissue from *Drosophila* was dissected after the exercise protocols. For the chronic exercise condition, flies were subjected to 7 days of exercise (2 hours per day) using the ClimbMaster apparatus, and muscle tissue was dissected immediately after the final exercise session. For the “acute exercise with insulin injection” condition, flies were subjected to a single 3-hour exercise session, followed by 1 hour of recovery before injection with hemolymph-like saline (HL3.1) or insulin in HL3.1. The HL3.1 saline solution contained 70 mM NaCl, 5 mM KCl, 1.5 mM CaCl2, 20 mM MgCl2, 10 mM NaHCO3, 115 mM sucrose (an inert osmolyte in this situation), and 5 mM trehalose to mimic the ionic composition of *Drosophila* hemolymph. Muscle tissue was dissected 4 hours post-injection to assess the effects of insulin. Samples in all cases included only the thorax with legs and wings. The gut and ventral nerve cord (VNC) were removed, and since little fat is present in the thorax, these preparations were largely composed of muscle tissue. For each condition, four to six biological replicates were collected, with each replicate consisting of thoracic muscle tissue from three flies. Total RNA was extracted using the NucleoSpin RNA kit (Macherey-Nagel, #740955), according to the manufacturer’s protocol. Tissue samples were homogenized in 2-mL Eppendorf tubes containing RA1 lysis buffer with 1% beta-mercaptoethanol, using a TissueLyser LT bead mill (Qiagen) and 5-mm stainless steel beads (Qiagen, #69989). Complementary DNA (cDNA) was synthesized using the High-Capacity cDNA Synthesis Kit (Applied Biosystems, #4368814), according to the manufacturer’s instructions. Real-time quantitative PCR (qPCR) reactions were performed using RealQ Plus 2x SYBR Green Master Mix (Ampliqon, #A324402) on a QuantStudio 5 system (Applied Biosystems). The housekeeping gene *Rp49* was used to normalize gene expression values across all samples. Primer sequences used for qPCR are listed in Supplementary Table 1.

### Metabolite and protein measurements

Whole-body levels of triacylglycerides (TAG), glycogen, and protein were measured in male and female flies following single-day or chronic exercise regimens. For the former group, flies were subjected to a single 3-hour exercise session, while the chronic exercise group underwent 1 hour of daily exercise for five consecutive days, followed by an additional 3-hour session on the final day. Fly samples were prepared by homogenizing 4 flies per sample in PBS containing 0.1% Tween-20 (Sigma, #1379) using a TissueLyser LT bead mill (Qiagen) and 5-mm stainless-steel beads. Homogenates were used to quantify TAG, glycogen, and protein levels. TAG levels were determined by hydrolyzing triglycerides into glycerol using Triglyceride Reagent (Randox TAG TR210). The liberated glycerol was then quantified using Free Glycerol Reagent (Sigma, #F6428) in a colorimetric assay. Glycogen levels were measured from the same extracts by hydrolyzing glycogen into glucose using amyloglucosidase (Sigma, #A7420), followed by a colorimetric glucose assay using Glucose Reagent (Sigma, #GAGO20). Protein levels were quantified using a bicinchoninic acid (BCA) assay. Extracts were combined with BCA solution (Sigma, #B9643) and copper sulfate solution (Sigma, #C2284), with bovine serum albumin (BSA) serving as the protein standard (Sigma, #P0914). The absorbance of each sample was measured in a 384-well plate at 500 nm for TAG assays, 540 nm for glycogen assays and at 562 nm for protein measurements using an EnSight multimode plate reader (PerkinElmer). Standard curves were generated to calculate TAG, glycogen, and protein concentrations. TAG and glycogen levels were normalized to protein concentrations to account for variability in the size and number of flies in each sample.

### Tissue lysis and immunoblotting

Samples for Western blot analysis of insulin pathway activation through phospho-AKT were prepared by dissecting tissues into ice-cold RIPA lysis buffer. The lysis buffer contained 50 mM Tris-HCl (pH 7.5), 150 mM NaCl, 0.5% sodium deoxycholate, 1% Nonidet P-40, and 0.1% SDS. Immediately before use, the buffer was supplemented with Roche Complete Protease Inhibitor Cocktail with EDTA (used at 4x, Roche #11836153001), PhosStop phosphatase inhibitor (used at 2x, Roche #PHOSS-RO), 11 mg/mL β-glycerophosphate (from a 110 mg/mL 10x stock), ∼100 mM NaF (from a saturated ∼1 M 10x stock in RIPA buffer), and Benzonase endonuclease (Sigma-Aldrich, #G9422). Six male *Drosophila* torsos (without heads and ventral nerve cords) were pooled per sample. Samples were kept on ice during dissection and until further processing. For protein denaturation, every two replicates were processed in order to prevent protein degradation and de-phosphorylation: the samples were mixed 1:1 with 2x Laemmli Sample Buffer (Bio-Rad #1620737) containing 5% 2-mercaptoethanol and homogenized using a TissueLyser bead mill (Qiagen) with a 5-mm steel bead. Homogenized samples were immediately heat-denatured at 95°C for 5 minutes. Following denaturation of all samples, samples were centrifuged at 14,000 rpm for 15 minutes at room temperature to pellet tissue debris. The supernatant was transferred to new Eppendorf tubes and centrifuged again under the same conditions to clarify the lysate further. The supernatant from the middle layer was collected into fresh tubes and stored at -80°C until use. Before loading onto SDS-PAGE gels, samples were thawed to room temperature and centrifuged again at 14,000 rpm for 5 minutes. Proteins were separated using precast 4-20% gradient Mini-Protean TGX polyacrylamide gels (Bio-Rad #4561094) in a BioRad Mini-PROTEAN Tetra Vertical Electrophoresis Cell system at 150 V for 30-45 minutes. The running buffer contained 3.02 g Tris base, 18.8 g glycine, and 10 g SDS per liter. PageRuler Plus Prestained Ladder (ThermoFisher Scientific #815-968-0747) was used as a molecular weight standard. Separated proteins were transferred to 0.2 µm polyvinylidene difluoride (PVDF) membranes using the Trans-Blot Turbo Transfer System (Bio-Rad #1704156). Membranes were cut at ∼35 kDa to probe for AKT (∼60 kDa) on the upper section and histone H3 (∼15 kDa) on the lower section. Membranes were blocked for 1 hour at room temperature in Intercept blocking buffer (TBS) (LI-COR #927-60001) with 0.2% Tween-20. Primary antibodies were added to the blocking buffer overnight at 4°C with gentle shaking: rabbit anti-phospho-AKT (Cell Signaling Technology #4054, 1:1000), rabbit anti-histone H3 (Abcam #1791, 1:5000), and rabbit anti-AKT (pan) (Cell Signaling Technology #4691S, 1:1000).Membranes were washed three times for 15 minutes in PBS with 0.1% Tween-20 at room temperature. Goat anti-rabbit IgG H+L HRP-conjugated secondary antibody (ThermoFisher #31466, 0.5mg/mL in 50% glycerol) was diluted 1:10,000 in Intercept blocking buffer with 0.2% Tween-20 for 90 minutes at room temperature. Chemiluminescence detection was performed using ECL Prime Western Blotting Detection Reagents (Amersham #RPN2232) and imaged with an Odyssey Fc gel reader (LI-COR). For reprobing, membranes were stripped of phospho-AKT antibodies using Alfa Aesar Stripping Buffer (#J60925) for 30 minutes at 37°C. Membranes were imaged to insure full stripping of antibodies before continuing with reprobing. Membranes were washed three times for 15 minutes in PBS with 0.1% Tween-20, re-blocked with Intercept blocking buffer (LI-COR #927-60001), and re-stained for total AKT(pan) following the procedure described above.

### Mass-spectrometric phosphoproteomics and bioinformatics

Dissected thorax/leg/wing samples were lysed by sonicating once for 30 seconds at 80% amplitude in a solution containing 5% sodium deoxycholate (SDC) and 50 mM HEPES (pH 8.5) supplemented with Roche cOmplete protease inhibitor and PhosSTOP phosphatase inhibitors (Sigma #04693132001 and #PHOSS-RO). Lysates were diluted 1:1 to a final concentration of 2.5% SDC and sonicated twice more for 30 seconds each at 60% amplitude. Proteins were then denatured at 99 °C for 10 minutes, and insoluble material was pelleted by centrifugation at 20,000×g for 20 minutes. Protein concentration was measured using the Pierce BCA Protein Assay (ThermoFisher Scientific). A total of 75 µg of protein was reduced with 10 mM DTT for 20 minutes, followed by alkylation with 20-mM iodoacetamide for another 20 minutes. Proteins were digested overnight at 37 °C with 5% trypsin, with an additional 1% trypsin added the following day, and digestion continued for a further 2 hours. An equivalent of 50 µg of peptides was labeled with 18-plex tandem mass tags (TMTpro) according to the manufacturer’s protocols. Labeled peptide samples were combined into one tube, and SDC was precipitated through acidification and pelleted by centrifugation at 20,000×g for 10 minutes. The supernatant was transferred to a fresh tube and vacuum-dried to a final volume of 150 µL. Phosphopeptides were enriched using TiO2 beads, and both phosphopeptides and non-modified peptides were fractionated using high-pH reversed-phase (RP) fractionation as previously described^47^. Fractions were vacuum-dried and re-solubilized in 3.5 µL of 0.1% formic acid (FA). The re-solubilized peptides were loaded on a 20-cm analytical column (100-µm inner diameter) packed with ReproSil-Pur C18 AQ 1.9-µm RP material using an EASY-nanoLC system coupled to an Orbitrap Eclipse Tribrid mass spectrometer. Peptides were eluted from the column using a gradient with decreasing solvent A (0.1% formic acid) and increasing solvent B (95% acetonitrile, 0.1% formic acid). The gradient began at 100% solvent A and transitioned to 75% solvent A and 25% solvent B over 77 minutes. Between 77 and 91 minutes, the gradient shifted to 60% solvent A and 40% solvent B, followed by a rapid increase to 95% solvent B by the 92nd minute. The mass spectrometer was operated in positive mode at Orbitrap resolution of 120,000 with a scan range (m/*z*) of 350-1600 and a cycle time of 2 seconds. Fragment ions were generated using higher-energy collision-induced dissociation (35%) and detected at the MS2 level with an isolation window of 0.7 Da, a resolution of 50,000, and a maximum injection time of 100 milliseconds for non-modified peptides or 200 milliseconds for phosphopeptides.

Raw output was searched against the *Drosophila* proteome FASTA file (version 11.12.2024) in Proteome Discoverer (PD) version 2.5.0.400 (ThermoFisher Scientific) using the SEQUEST HT search engine, with a maximum of 2 missed cleavages allowed. Static modifications used included TMTpro on N-terminus, TMTpro on K, and carbamidomethyl on C. Dynamic modifications included oxidation on M, deamination on N, and phosphorylation on S, T or Y. Data analysis was performed using Perseus (version 2.0.10.0) and Jupyter Notebook (Python version 3.11.5). Phosphorylated peptides were filtered, and for statistical significance was assessed using a two-way ANOVA with between exercise (exercise vs. no exercise) and treatment (HL3.1 vs. insulin). False discovery rate (FDR) correction was applied using the Benjamini-Hochberg (BH) procedure, with significance thresholds set at FDR<0.2. For the experiment in which animals were subjected to exercise or no exercise without injection (Fig. 8e and 8g), statistical significance was determined using Student’s *t*-test with permutation-based FDR control, also set at FDR<0.2. The percentage of phosphorylated S, Y, or T residues was calculated by dividing the number of phosphorylated modifications for each residue type by the total number of phosphorylated peptides. Principal-component analysis (PCA) was conducted on z-scored data using the PCA implementation from the *scikit-learn* Python library. To further illustrate group clustering, 50% probability contours were added around the groups using kernel density estimation (KDE). Statistically significant z-scored proteins were clustered into two groups using the KMeans algorithm from *scikit-learn* (k=2), identifying major protein groupings based on abundance profiles. To reorder samples (columns), hierarchical clustering was applied using pairwise Euclidean distances and Ward’s linkage method via the linkage function from the *SciPy* library. Heatmaps visualizing protein clusters and sample relationships were generated using the *Seaborn* library.

## Acknowledgements

We thank the Technical Support Group, Mechanical Department, Niels Bohr Institute, University of Copenhagen, for their assistance in developing the ClimbMaster hardware and control software. This work was supported by grants from the Novo Nordisk Foundation to KR (NNF22OC0078344 and NNF19OC0054632). The Zeiss LSM-900 confocal microscope and the PerkinElmer EnSight plate reader were funded through generous grants from the Carlsberg Foundation (CF19-0353 and CF17-0615, respectively) awarded to KR.

## Author contributions

AM, MJT, and KR conceived and designed the study. AM, OK, ML, AHS, IB, MRL, NA and MJT performed the experiments and analyzed the data. AM, MJT, and KR wrote the manuscript.

