## Supplemental table 1 for "High-throughput assessment of exercise-induced adaptations and muscle function in health and ageing"

### Supplementary Table S1: Oligo sequences used for qPCR

| Oligos | Forward | Reverse |
|---|---|---|
| *Rp49* | AGTATCTGATGCCCAACATCG | CAATCTCCTTGCGCTTCTTG |
| *InR* | CTCAGCCATACCAGGGACTTT | CTCTCCATAACACCGCCATC |
| *4EBP* | TGCCCATGATCACCAGGAAG | TCGTAGATAAGTTTGGTGCCTCC |
| *Pepck* | TCAATGGCGAATCCTGCTAC | TCCTTCACGTCCACCTTATCC |
